# Ecological factors shaping the ectoparasite community assembly of the Azara’s Grass Mouse, *Akodon azarae* (Rodentia: Cricetidae)

**DOI:** 10.1101/2023.05.17.541149

**Authors:** Colombo Valeria Carolina, Lareschi Marcela, Monje Lucas Daniel, Antoniazzi Leandro Raúl, Morand Serge, Beldomenico Pablo Martín

**Affiliations:** Laboratorio de Ecología de Enfermedades, Instituto de Ciencias Veterinarias del Litoral (ICIVET-Litoral), Universidad Nacional del Litoral (UNL) / Consejo Nacional de Investigaciones Científicas y Técnicas (CONICET), R.P. Kreder 2805, 3080, Esperanza, Argentina; Evolutionary Ecology Group, Department of Biology, University of Antwerp, 2610, Wilrijk, Belgium; Servicio de Neurovirosis, INEI-ANLIS “Dr. Carlos G. Malbrán”, Av. Vélez Sarsfield 563, C1282AFF, Ciudad Autónoma de Buenos Aires, Argentina; Centro de Estudios Parasitológicos y de Vectores (CEPAVE) (CONICET-UNLP), Bv. 120 s/n e/ 60 y 61, 1900, La Plata, Argentina; Instituto de Bio y Geociencias del NOA (CONICET), 9 de julio 14, 4405, Rosario de Lerma, Argentina; Maladies infectieuses et vecteurs: écologie, génétique, évolution et contrôle (MIVEGEC), Université de Montpellier, CNRS, IRD, 34090, Montpellier, France

## Abstract

Parasites are integral members of the global biodiversity. They are useful indicators of environmental stress, food web structure and diversity. Ectoparasites have the potential to transmit vector-borne diseases of public health and veterinary importance and to play an important role in the regulation and evolution of host populations. The interlinkages between hosts, parasites and the environment are complex and challenging to study, leading to controversial results. Most previous studies have been focused on one or two parasite groups, while host are often co-infected by different taxa. The present study aims to assess the influence of environmental and host traits on the entire ectoparasite community composition of the rodent Akodon azarae. A total of 278 rodents were examined and mites (Mesostigmata), lice (Phthiraptera), ticks (Ixodida) and fleas (Siphonaptera) were determined. A Multi Correspondence Analyses was performed in order to analyse interactions within the ectoparasite community and the influence of environmental and host variables on this assembly. We found that environmental variables have a stronger influence on the composition of the ectoparasite community of A. azarae than the host variables analysed. Minimum temperature was the most influential variable among the studied. In addition, we found evidence of agonistic and antagonistic interactions between ticks and mites, lice and fleas. The present study supports the hypothesis that minimum temperature play a major role in the dynamics that shape the ectoparasite community of A. azarae, probably through both direct and indirect processes. This finding becomes particularly relevant in a climate change scenario.

## Introduction

Parasites are integral members of the global biodiversity. They are useful indicators of environmental stress, food web structure and biodiversity (Marcogliese 2005) and play an important role in the regulation and evolution of host populations and communities (e.g., Morand and Poulin 1998; Morand and Bordes 2015). Parasite communities are composed of and aggregated in their hosts heterogeneously as a result of differences on the chances of hostparasite encounters and host susceptibility (Combes 1991; Johnson and Hoverman 2014; Gourbiere et al. 2015). One of the phenomena that drive these asymmetries is the difference on the characteristics of their life-cycle. In particular for ectoparasites, their communities are composed by different groups and taxa, whose life cycle characteristics can be widely different. While some ectoparasites develop their entire life cycle on their hosts (e.g. lice), others spend most of their time in the environment (e.g. ticks), where they deposit eggs and/or immatures molt or grow up (e.g. fleas), and contact a host only during a short proportion of their life for food and/or mating purposes. The feeding ecology can also be quite different, ranging from hematophagous diets through living at the expense of skin detritus. Based on these differences, we can expect that host and environmental characteristics will asymmetrically bias the presence and abundance of the different ectoparasites; highlighting the need to consider the network of interactions (both direct and indirect) that occurs between parasites, hosts and the environment to obtain a mechanistic understanding of parasite communities (Pedersen and Fenton 2006). In addition, the habitat of hosts and ectoparasites can be modified by human land use and climate change, altering hosts’ diet, distribution and diversity (e.g., Skov and Svenning 2004; Sillero and Carretero 2013; Werner and Nunn 2020). Thus, the survival, chances of encounter and establishment of ectoparasites in susceptible hosts can be altered (Hieronimo et al., 2014; Shilereyo et al. 2022), potentially reflected as changes in their community composition. The interlinkages between the different components of these ecological networks are complex and challenging to study, leading to controversial results (Estrada Peña et al. 2004; Kiffner et al. 2011). Most of these studies have been focused on one or two parasite groups (e.g., Estrada Peña et al. 2004; Kiffner et al. 2011; Hieronimo et al. 2014; Shilereyo et al. 2022), while host are often co-infected by different taxa simultaneously (Bordes and Morand 2011). As a result, the mechanisms that shape within-host parasite communities remain unclear.

Furthermore, ectoparasites can transmit vector-borne diseases of public health and veterinary importance, especially parasites of rodent hosts, which are among the main reservoirs of zoonotic diseases worldwide (Han et al. 2015). In South America, the Azara’s Grass Mouse, *Akodon azarae* (Rodentia: Cricetidae: Sigmodontinae), is one of the most abundant native rodent species distributed from Rio Grande do Sul in southern Brazil, and eastern Paraguay through Uruguay to central Argentina (Bilenca and Kravetz, 1998; Patton et al. 2015). It is considered a reservoir and/or amplifier host of multiple infectious agents, such as *Bartonella* spp., *Leptospira interrogans,* Orthohantavirus, and *Rickettsia parkeri* (Levis et al. 1998; Colombo 2016; Colombo et al. 2018; Schott et al. 2020), as well as host of several ectoparasite species (e.g., Navone et al. 2009; Nava et al. 2011; Colombo 2016; Lareschi 2020). How different traits can shape the ectoparasite communities in their rodent hosts provides the basis of the understanding of host – parasite – environment dynamics with diverse applications in community and disease ecology.

The aim of the present study was to assess the influence of environmental and host traits on the ectoparasite community assembly, composed of mites (Mesostigmata), lice (Phthiraptera), ticks (Ixodida) and fleas (Siphonaptera), of the rodent host *A. azar*a*e* in a South American location.

## Materials and methods

### Study area

The study was conducted at the Estación Experimental Agropecuaria (EEA) Delta, Instituto Nacional de Tecnología Agropecuaria (INTA), Campana (34°11 S, 58°50 W), Buenos Aires, Argentina (Fig. 1). The site is located in the lower Parana River Delta region, which is the southern extension of the Paranense Province of the Amazonic Phytogeo-graphic Dominion (Cabrera 1994). Besides the native vegetation, the site has areas with commercial forestations of *Populus* spp. and *Salix* spp. The site is characterized by levees that surround dry areas as well as temporarily or permanently flooded marshes. In addition, in the study area there is a herd of beef cattle consisting of twenty-one Aberdeen Angus cows maintained at a density of approximately one cow per hectare. The climate is temperate with a mean annual temperature of 16.7°C and a mean annual rainfall of 1000 mm with an undefined rainy season (Kandus and Malvárez 2004).

**Fig. 1.**
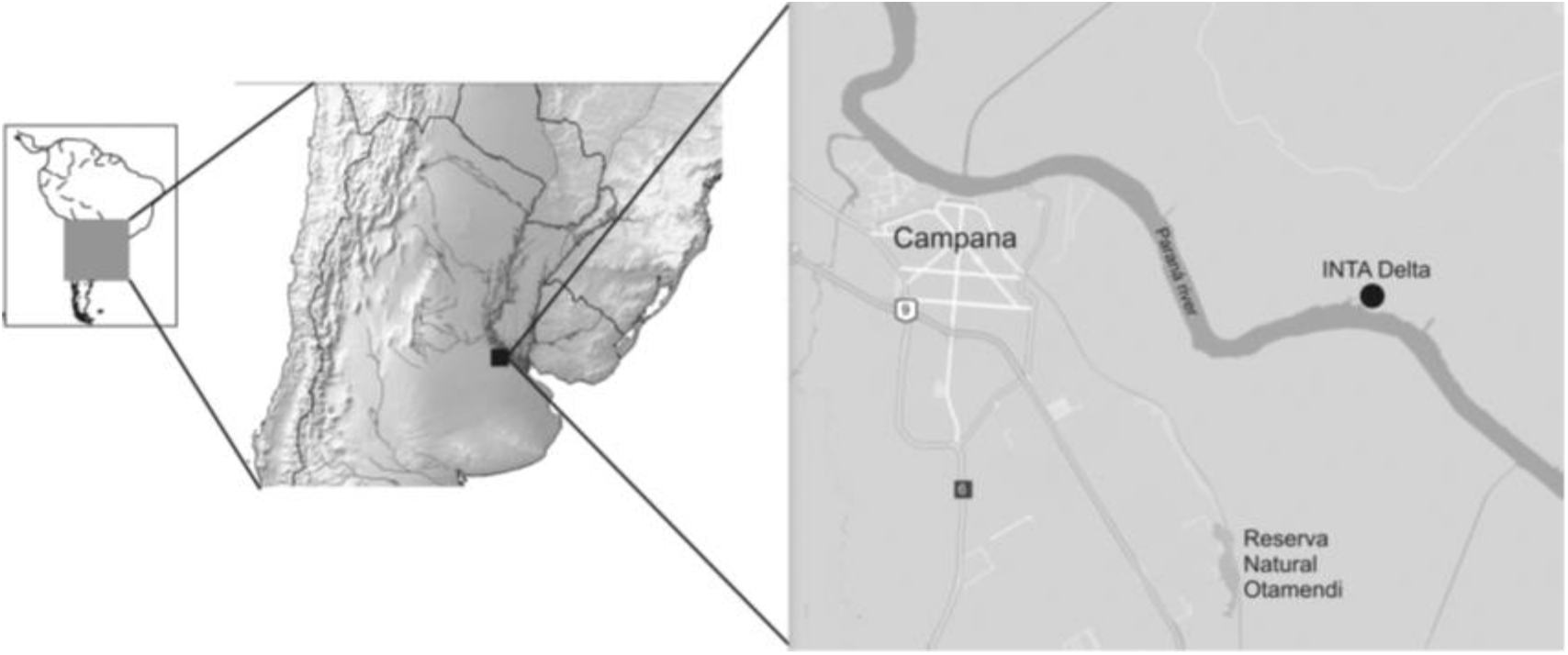
Map showing the location of the study area and the sampling plots, INTA EEA Delta (34°11 S, 58°50 W), Campana, Buenos Aires province, Argentina.

### Data collection

Rodents were captured from November 2010 through October 2012 in 3-night trapping sessions carried out every 5 weeks. Four trapping grids were set out at 4 different sites, each grid was 200 m apart from each other and consisted of squares with 12 Sherman-type live-traps in the corners and 2 Ugglan-type live-traps in the middle of the square, baited with pelleted dog food. Within a site, two of the grids were located in places with natural grassland and the other two with commercial forestations. Half of the sites were located in extensive cattle raising lands and the other half in areas where cattle was absent. Every morning trapped rodents were transported to a field lab, anesthetized by inhalation of Isoflurane, sacrificed by cervical dislocation and then conserved in individual plastic bags with ethanol 96°. Blood samples were taken by heart puncture and collected in heparin-coated capillary tubes and Eppendorf tubes without anticoagulant. Rodents were later identified to the species level by assessing cranium morphology. Although other rodent species (Cricetidae: Sigmodontinae) were trapped during this study (namely, *Oxymycterus rufus, Oligoryzomys flavescens, Oligoryzomys nigripes, Scapteromys aquaticus* and *Deltamys kempi*, as described by Colombo et al. (2013), only *A. azarae* was the abundant enough to carry out the multivariable analyses desired.

### Ectoparasites

Each rodent was examined in the laboratory with magnifying lens and all ectoparasites were collected and counted. Ticks were determined following Estrada Peña et al. (2005), Martins et al. (2010) and Marques et al. (2004). The remaining ectoparasites were identified under light microscope. Mites were cleared in lactophenol and mounted in Hoyeŕs medium; fleas and lice were cleared by using 10% KOH and mounted in Canadian balsam. Strandtmann & Wharton (1958) and Furman (1972) and Lareschi (2020) were followed for mites identification; Johnson (1957), Smit (1987) and Linardi and Guimarães (2000) for flea; and Johnson (1972) and Durden and Musser (1994) for lice. Ectoparasites were also compared with specimens deposited in different collections of Argentina. All ectoparasites collected were determined to the species level except for Polygenis fleas that were determined to the subgenera level.

### Statistical analysis

Data analyses were conducted in two steps. First, in order to reduce ectoparasite, host and environmental variables superfluous information, we performed Principal Components Analyses (PCA) with (1) *A. azarae* ectoparasite abundances including the ticks *Amblyomma triste* and *Ixodes loricatus* (Acari: Ixodidae), the mites *Androlaelaps fahrenholzi*, *Androlaelaps azarae* and *Ornithonyssus bacoti* (Acari: Mesostigmata), the fleas *Polygenis* subgenera *Polygenis* and *Neopolygenis*, and *Craneopsylla minerva minerva* (Insecta: Siphonaptera), and the louse *Hoplopleura aitkeni* (Insecta: Phthiraptera). (2) *Host parameters*: body length (cm); weight (grams); age (juvenile, sub-adult and adult); sex (female and male); body condition score (1 through 10); and reproductive status (active/inactive). (3) *Environmental variables*: cattle (present or absent); type of vegetation (natural grassland or implanted forest); season (summer, autumn, winter and spring as determined by solstices and equinoxes) and minimum temperature (Tmin) (C°) measured as a weekly average at different time lags (Tmin-0 through Tmin-8) and as an unique measure of the trapping session day (Tmin-pres).

According to the PCAs, the flea subgenera *Polygenis* and *Neopolygenis* showed high correlation values, thus they were grouped into *Polygenis* spp. for further analyses. The host variables age and weight were excluded for further analyses as they showed high correlation between them and with reproductive status and body length. Among the environmental variables, most of the time lags for Tmin showed high correlation between them and with the variable season; therefore we selected Tmin-pres and Tmin-8 for further analyses. These variables did not correlate with each other and could represent the minimum temperature of the date of the trapping session and up to 4 weeks beforehand (Tmin-pres) and between week 5 and 8 before the sampling date (Tmin-8).

Second, in order to analyse the association of environmental and host factors with the entire ectoparasite community composition (parasite prevalence) including ticks, mites, fleas and lice, and the presence of interactions within the ectoparasite community, a Multi Correspondence Analysis (MCA) was performed. MCA is a nonlinear multivariate method that allows the inclusion of all ectoparasite taxa in the same analyses, overcoming the limitations of regression models. Only ectoparasites with prevalence values equal or higher than 10% were considered including *A. triste* and *I. loricatus*, *A. fahrenholzi*, *An. azarae, Polygenis* spp., *C. minerva minerva*, and *H. aitkeni*. The host factors included as additional variables were body length, sex, body condition score, and reproductive status. The environmental factors included were presence/absence of cattle, type of vegetation, Tmin-pres, and Tmin-8. Both PCA and MCA were conducted using the *FactoMinR* package for the analyses and *factoextra* for data visualization with the statistical software R (R Foundation for Statistical Computing, http://www.r-project.org).

## Results

In the 278 rodents analysed, the mite *An. azarae* was the most prevalent species of ectoparasite collected (76.2%), followed by the louse *H. aitkeni* (37%), *A. triste* ticks and *Polygenis* spp. fleas (33%). Less than a third of the rodents had *A. fahrenholzi* mites (27.6%), *C. minerva minerva* fleas (24%) and *I. loricatus* ticks (19.5%). Finally, *O. bacoti* mites were the less prevalent ectoparasite (4.6%). Regarding ectoparasite abundance, the louse *H. aitkeni,* the mite *A. azarae*, the tick *A. triste* and *Polygenis* spp. fleas were the four most abundant ectoparasites in decreasing order.

Regarding the MCA, the first three dimensions accounted for 53.7% of the statistical inertia. The dimension 1, explaining 20.7% of the variation in the ectoparasite community, primarily described the occurrence of the flea *C. minerva minerva* (R2=40%, P<0.001) and the mite *A. fahrenholzi* (R2=30%, P<0.001) (Fig. 2 and 5). The mite *An. azarae* (R2=23%, P<0.001) and the ticks *I. loricatus* (R2=21%, P<0.001) and *A. triste* (R2=20%, P<0.001) showed a lower contribution to the total variance of the dimension (Fig. 2 and 5). The two tick species studied and the mite *A. fahrenholzi* were clustered close together (Fig. 3a) suggesting a co-occurrence pattern. The dynamics observed in this dimension were ruled mostly by the environmental variable Tmin-8 (Corr=21%, P=0.0003), weakly followed by the host variable body length (Corr=14%, P=0.01) (Fig. 3b).

**Fig. 2.**
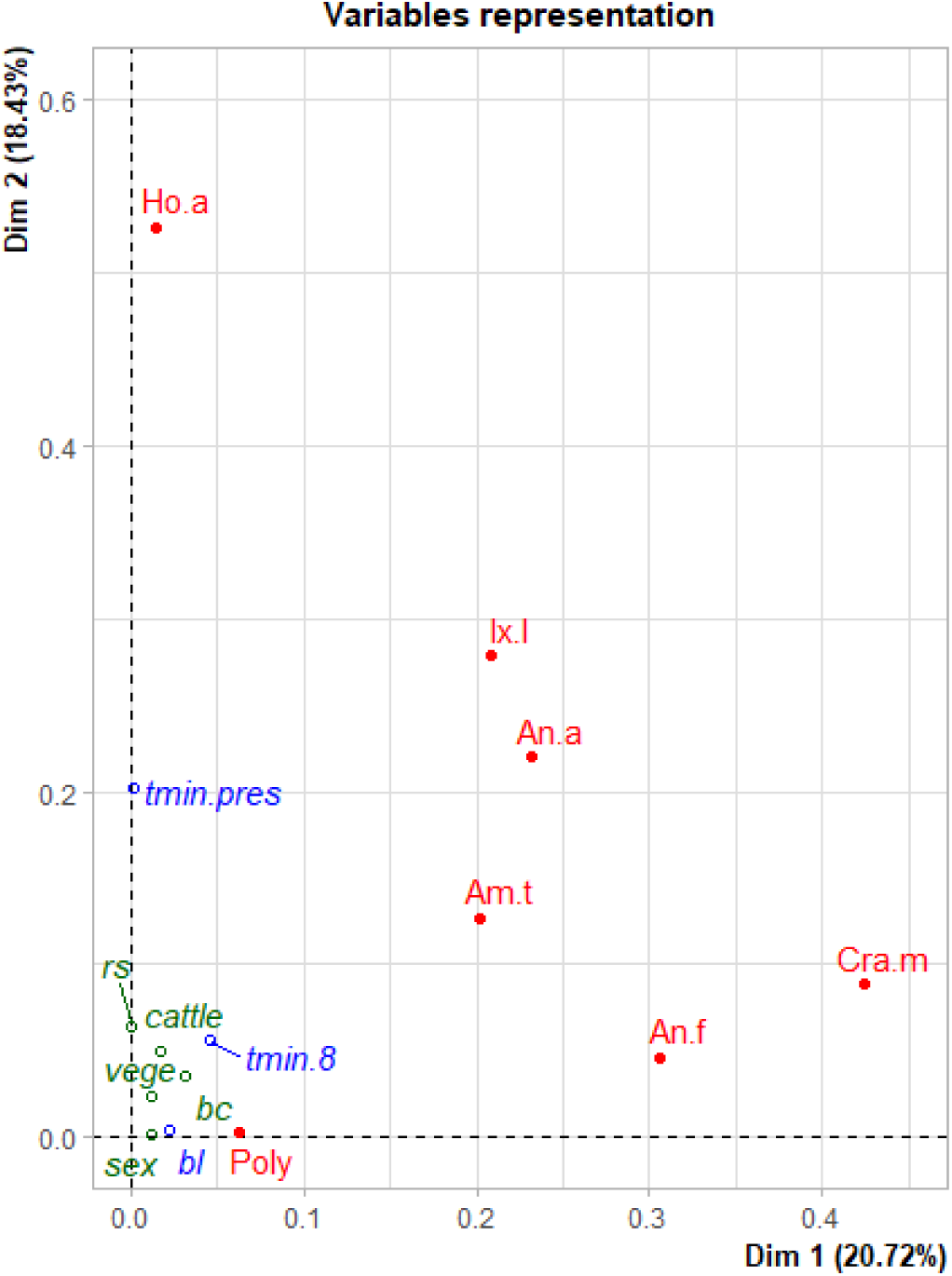
Graphical representation of the results of the multiple correspondence analysis (MCA). Variables distribution in dimensions 1 and 2. Ectoparasites (An.a: *An. azarae*, An.f: *A. fahrenholzi*, Am.t: *A. triste*, Cra.m: *C. Minerva minerva*, Ho.a: *H. aitkeni*, Ix.l: *I. loricatus*, Poly: *Polygenis* spp.); host parameters (bc: body condition, bl: body length, rs: reproductive status, sex: host gender) and environmental variables (cattle: presence/absence of cattle, tmin.pres: minimum temperature of the month of sampling, tmin.8: minimum temperature from weeks 5 to 8 before sampling, vege: natural grassland/implanted forest).

**Fig. 3.**
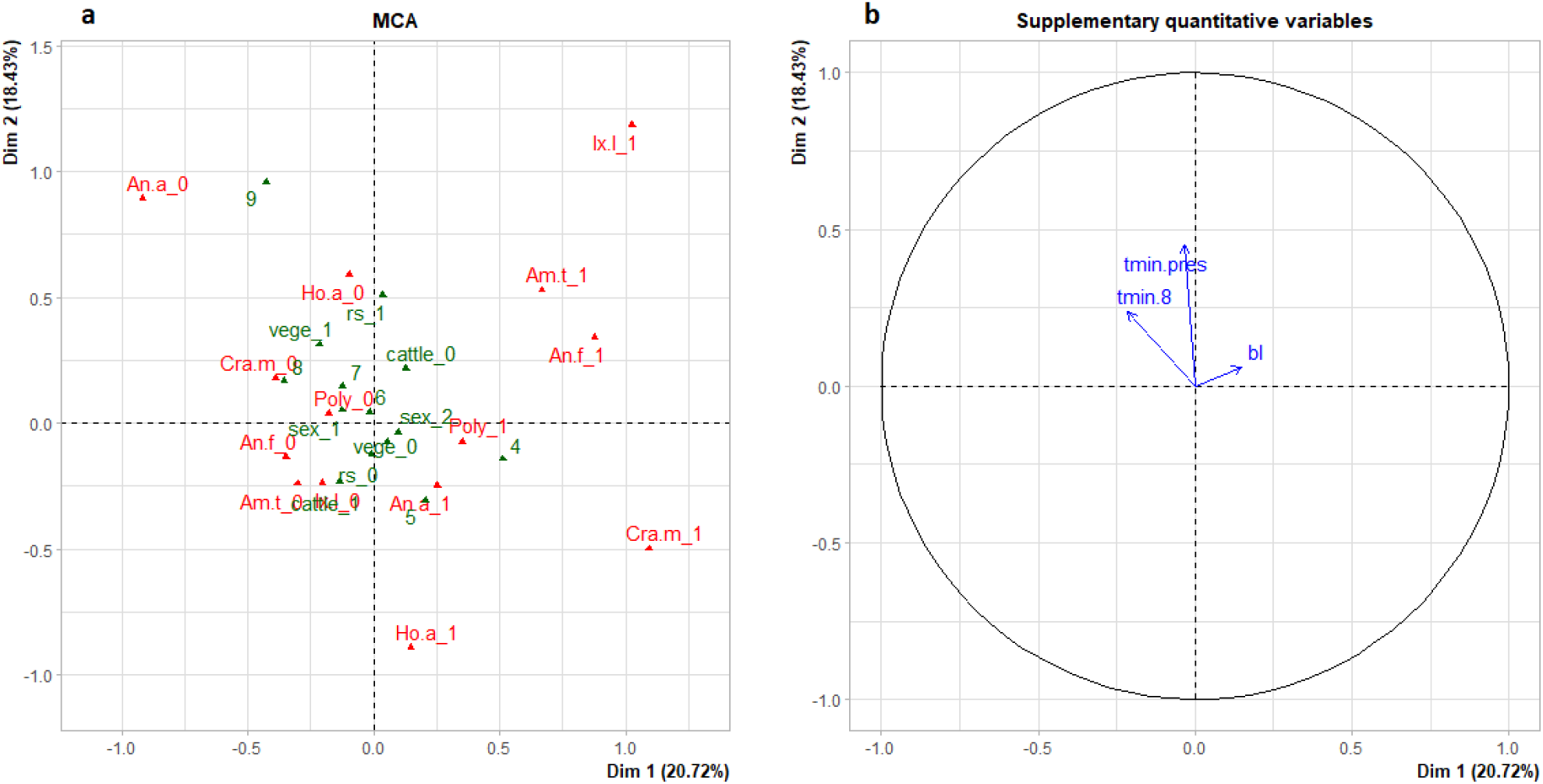
Graphical representation of the results of the multiple correspondence analysis (MCA). (a) Variables distribution in dimensions 1 and 2. Ectoparasites presence (1)/absence (0) (An.a: *An. azarae*, An.f: *A. fahrenholzi*, Am.t: *A. triste*, Cra.m: *C. Minerva minerva*, Ho.a: *H. aitkeni*, Ix.l: *I. loricatus*, Poly: *Polygenis* spp.); qualitative host variables (bc: body condition from 5 to 9), rs: reproductive status active(1)/inactive(0), sex: female(1), male(2)) and qualitative environmental variables (cattle: presence(1)/absence(0), vege: natural grassland(0)/implanted forest(1)). (b) Quantitative variables representation (bl: Body length, tmin.pres: minimum temperature of the month of sampling, tmin.8: minimum temperature from weeks 5 to 8 before sampling)

The second dimension, explaining 18.4% of the variation, was characterized mostly by the parasitism of the louse *H. aitkeni* (R2=52%, P<0.001), followed by *I. loricatus* (R2=27%, P<0.001), and *An. azarae* (R2=22%, P<0.001) (Fig. 2 and 5). *Hoplopleura aitkeni* and *I. loricatus* showed a negative interaction pattern (Fig. 3a). These ectoparasite’s dynamics were mainly explained by the environmental factors Tmin-pres (P<0.001), explaining the 45% of the variability of the dimension, and secondarily by Tmin-8 (Corr=23%, P<0.001) (Fig. 3b).

Finally, the third dimension explained the 14.6% of the variation and was characterized by the parasitism of the flea *Polygenis* spp (R2=60%, P<0.001), followed by the co-occurrence with the tick *A. triste* (R2=20%, P<0.001) (Fig. 4 and 5). These associations were mostly explained by the environmental variable Tmin-pres (Corr=22%, P<0.001).

**Fig. 4.**
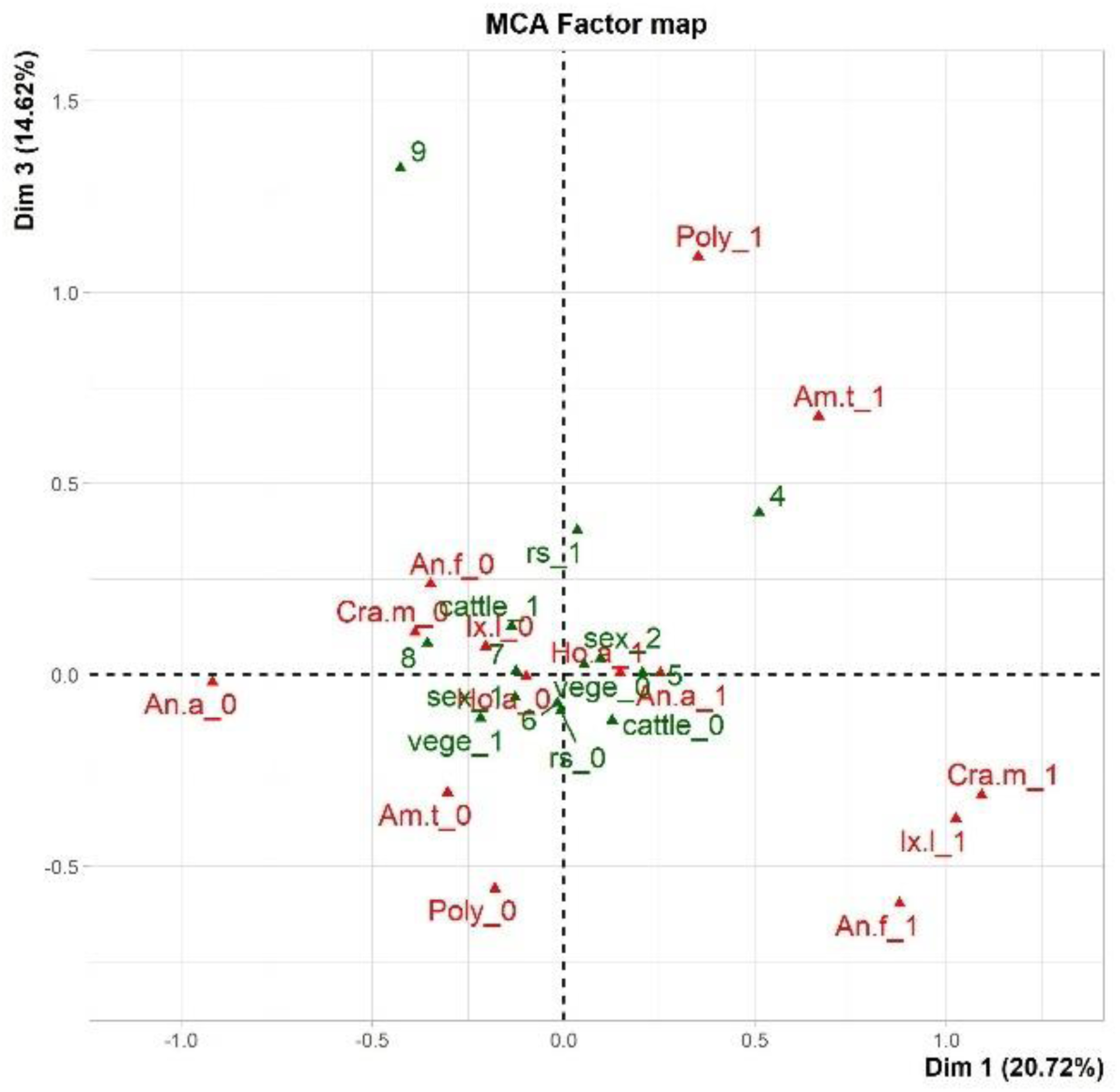
Graphical representation of the results of the multiple correspondence analysis (MCA). Variables distribution in dimensions 1 and 3. Ectoparasites presence (1)/absence (0) (An.a: *An. azarae*, An.f: *A. fahrenholzi*, Am.t: *A. triste*, Cra.m: *C. Minerva minerva*, Ho.a: *H. aitkeni*, Ix.l: *I. loricatus*, Poly: *Polygenis* spp.); qualitative host variables (bc: body condition from 5 to 9), rs: reproductive status active(1)/inactive(0), sex: female(1), male(2)) and qualitative environmental variables (cattle: presence(1)/absence(0), vege: natural grassland(0)/implanted forest(1)).

**Fig 5.**
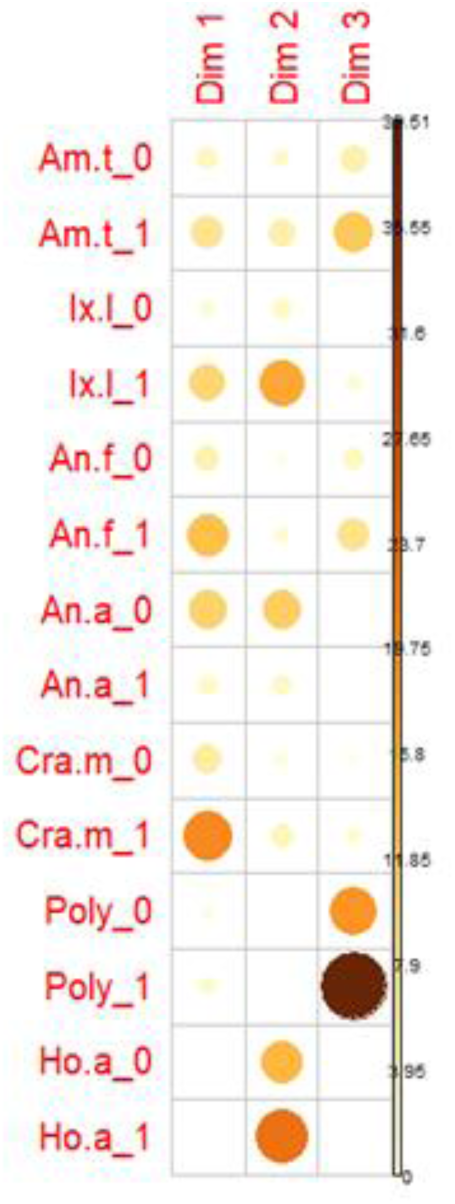
Contributions of each ectoparasite (0: absence, 1:presence) to the definition of the dimensions 1, 2 and 3 of the MCA. (An.a: *An. azarae*, An.f: *A. fahrenholzi*, Am.t: *A. triste*, Cra.m: *C. Minerva minerva*, Ho.a: *H. aitkeni*, Ix.l: *I. loricatus*, Poly: *Polygenis* spp.)

No additional associations with sufficient variance for statistical tests were clearly suggested by the MCA analysis.

## Discussion

The dynamics of the interlinkages between the ectoparasite community, the host and the environment are complex and difficult to unravel. In the present study, we found that environmental variables have a stronger influence on the composition of the ectoparasite community of *A. azarae* than the host variables analysed.

Environmental factors can bias the survival, behaviour, abundance and host encounter of ectoparasites during their off-host period. Three-host ticks, as *A. triste* and *I. loricatus*, spend most of their lifetime in the environment during oviposition, moulting and host questing. It is well-known that their dynamics are influenced by the saturation deficit, temperature, and relative humidity at the microclimate scale (Randolph and Storey, 1999; Randolph et al. 2002; Yoder et al. 2008; Raghavan et al. 2016). Temperature has been found to be an influential variable in the distribution, survival, behaviour and therefore density of questing ticks (e.g., Perret et al. 2000; Cumming et al. 2002; Estrada Peña et al. 2004), which will be reflected in their parasitism on hosts (Brunner and Ostfeld 2008). Unlike ticks, fleas and mites spend much of their environmental phase in host nests or burrows, with more protection from environmental changes. However, it is known that eggs and/or immature stages of fleas and some mites (e.g., *O. bacoti* and *A. fahrenholzi*) are influenced by temperature and humidity, affecting their survival and developing time, rate of oviposition in adults, feeding success and abundance, among others (Strandtmann and Wharton 1958; Radovsky 1985; Krasnov et al. 2001a, b; Krasnov 2008; Krasnov et al. 2007; Walter and Proctor 2013). Lice are considered permanent parasites as they spend their entire life on the host (Marshall 1981). However, this does not prevent their parasitism from being affected by environmental conditions. The effect of variables such as season, habitat, locality and host density on lice parasitism has been reported in several studies (e.g., Mize et al. 2011; Fernandes et al. 2012; Archer et al. 2014; Stanko et al. 2015). Warmer temperatures have been associated with higher occurrences of louse infestation in rodent hosts, as well as to the survival and development of pre-imaginal lice and the rate of oviposition of adults (Schrader et al. 2008; Colwell 2014; Stanko 2015). Environmental factors can also influence the diversity of ectoparasites. Liljesthröm and Lareschi (2018) found that environmental stress is a good predictor for ectoparasite species richness through its direct effect on the ectoparasites during their off-host period, an effect on the vegetation that provides host shelters or parasite nests, and a direct effect on rodent hosts.

Krasnov et al. (2004) found that flea species richness is affected little by parameters of the host and to a greater extent by parameters related to the host environment. The diversity of mites has also been strongly correlated with environmental factors; the taxonomic diversity of mites in their hosts can decrease with an increase in winter precipitation and/or air temperature (Krasnov et al. 2007). These changes in the diversity are then reflected in the ectoparasite community composition. In the present study, we found that the minimum temperature was the most influential variable in the ectoparasite community dynamics of the host *A. azarae*, regardless of their life-cycle characteristics.

Through environmental fluctuations, some ectoparasite species would be more prone to suffer consequences on their dynamics than others (Marshall 1981; Krasnov et al. 2022). These consequences can compromise not only the stability of a particular species but also that of the entire ectoparasite community due to interspecific interactions at the infracommunity level (Pedersen and Fenton 2006; Lutermann et al. 2015; Hoffmann et al. 2016; Krasnov et al. 2020). The occurrence and burden of flea, mite and tick species have been found to be associated with the parasitism of other ectoparasites as a result of agonistic or antagonistic interactions within their host (e.g., Krasnov et al. 2010; Colombo et al. 2015a, b; Krasnov et al. 2021). In this regard, environmental factors may not only have a direct effect on parasite species dynamics but also an indirect effect through interspecific interactions. In the present study, we found trends of agonistic and antagonistic interactions between ticks and mites, lice and fleas biased by minimum temperature conditions.

Furthermore, environmental factors can bias host density and behaviour (e.g., Hansson 1979; Kausrud et al. 2008; Sassi et al. 2015; Nater et al. 2018), indirectly influencing the ectoparasite community dynamics (Krasnov et al. 2002; Stanko et al. 2015; Krasnov et al. 2022). Krasnov et al. (2004) found that rodents from temperate regions had richer flea assemblages, probably due to deeper burrows in these regions. The level of rodent sociality is also considered a factor associated with the species richness and occurrence of directly transmitted ectoparasites (Bordes et al. 2007; Stanko et al. 2015). Host mating system, as well as host density, are considered factors that bias ectoparasite distribution (Brunner et al. 2008; Stanko et al. 2015; Wang et al. 2015; Elias et al. 2020), and can be explained through the number of direct contacts between hosts, as well as indirect contacts via visiting each other’s burrows; which facilitates an exchange of ectoparasites between individuals (see in Krasnov et al. 2022). In this sense, *A. azarae* has a polygynous mating system with a long breeding season (about 8 months), it uses shallow holes for nesting and occasionally dig burrows (see in Patton et al., 2015). These observations demonstrate how environmental factors can influence the composition of ectoparasites infracommunities through different ecological mechanisms. Changes in minimum temperature could modify the behaviour of *A. azarae*, indirectly influencing the ectoparasite community assembly.

Host traits have also been found to be related to ectoparasite parasitism. Host characteristics such as body length (e.g., Krasnov et al. 2004; Cardon et al. 2011; Fernandes et al. 2015), age and body mass (e.g., Soliman et al. 2001; Krasnov et al. 2006; Kiffner et al. 2011; Buchholz et al. 2017), and sex (Soliman et al. 2001; Krasnov et al. 2012; Fernandes et al. 2012) have been linked with the occurrence, burden, and/or diversity of different ectoparasite taxa. In the present study, the MCA showed little support for an association between the host variables studied and *A. azarae* ectoparasites. Body length was the only host variable slightly significant in the dimension that mostly describes the occurrence of the flea *C. minerva minerva*. This variable is an indicator of rodent age (Zullinger et al. 1984), which is highly correlated with the maturity of the immune system, the chances of host-parasite encounter, energy expenditure and compensation mechanisms within host-parasite interactions (Hudson et al. 2002; Hawlena et al. 2006; Garrido et al. 2016; Clerc et al. 2019). Krasnov et al. (2006) found that mean abundance and species richness of fleas increased with an increase of host age, but the patterns of change in flea aggregation and prevalence with host age were different depending on the rodent species studied. Furthermore, differences on the body surface of a host have been previously considered as a difference in the “niche offer” able to support a different number and diversity of parasites (Soliman et al. 2001; Korallo et al. 2007; Cardon et al. 2011; Fernandes et al. 2015), which is then reflected in the ectoparasite community composition. Our study supports the theory that body length is one of the factors in the intricate network of variables that shape the ectoparasite community, although not as relevant as minimum temperature.

In conclusion, our results support the hypothesis that minimum temperature play a major role in the dynamics that shape the ectoparasite community of *A. azarae*, probably through both direct and indirect processes. On the other hand, body length was the only host variable that showed some impact on the ectoparasite community assembly of *A. azarae*. These results demonstrate how influential changes in minimum temperature can be on the dynamics of an ectoparasite community, with potential consequences on their host ecology, as well as on the ecology of the pathogens they transmit. This finding becomes particularly relevant in a climate change scenario.

## Acknowledgements

Special thanks to Instituto Nacional de Tecnología Agropecuaria (INTA) EEA Delta del Paraná, Facultad de Ciencias Veterinarias (UNL), Santiago Nava, Alberto A. Guglielmone, Ulyses F.J. Pardiñas, Natalia Fracassi, Gerardo Mujica, and Cristian Ortiz

## Funding

This work was funded by Agencia Nacional de Promoción Científica y Tecnológica (PICT 2008-00090) and by Universidad Nacional del Litoral (CAI+D 2011/501 201101 00498).

Ethics approval: All procedures were carried out following the Directive 2010/63/EU and under the approval of the Ethic and Biosafety Committee of the Facultad de Ciencias Veterinarias, Universidad Nacional del Litoral, Argentina, the Dirección de Flora y Fauna de la Provincia de Buenos Aires and the Ministerio de Asuntos Agrarios de la provincia de Buenos Aires (N° 22500-17229/12, Disposición N° 48/12).

## Contributions

Conceptualization: Valeria Carolina Colombo, Serge Morand; Methodology: Valeria Carolina Colombo, Marcela Lareschi, Serge Morand, Pablo Martin Beldomenico, Lucas Daniel Monje, Leandro Raul Antoniazzi; Formal analysis and investigation: Valeria Carolina Colombo, Marcela Lareschi, Serege Morand, Pablo Martin Beldomenico; Writing – original draft preparation: Valeria Carolina Colombo, Marcela Lareschi; Writing – review and editing: Lucas Daniel Monje, Leandro Raul Antoniazzi, Serge Morand, Pablo Martín Beldomenico; Funding acquisition: Pablo Martín Beldomenico, Lucas Daniel Monje; Supervision: Marcela Lareschi, Serge Morand.

